# Elephant poaching continues in northern Botswana: spatial patterns and policy implications

**DOI:** 10.64898/2026.11.20.768173

**Authors:** Scott Schlossberg, Michael J. Chase

## Abstract

Illegal killing for ivory is the primary threat to the endangered African savanna elephant (*Loxodonta africana*). Botswana has the world’s largest elephant population and experienced little poaching during the continent-wide poaching outbreak of the early 2000’s. In 2018, however, evidence of widespread elephant poaching in Botswana began to appear, with an estimated 385 elephants poached. Here, we update the status of elephant poaching in northern Botswana with new data collected in 2023-2026. We also used spatial models to determine the factors associated with elephant poaching locations and to better understand poacher behavior. From late 2023 through mid-2026, we located 202 poached elephants in a ∼10,000 km^2^ area of northern Botswana, north and east of the Okavango Delta and mostly west of Chobe National Park. All poached elephants were believed to be adult males. Using point process models, we found that, depending on the method used to locate carcasses, poaching locations were associated with greater distance from roads, higher slope, lower distance to military posts, higher Enhanced Vegetation Index in the dry season, and greater distance from the Namibia border than random points. The poaching outbreak documented in 2018 was not transient: illegal killing for ivory appears to have continued. Because poachers are targeting adult males, their numbers could become depleted in our study area, which has implications for elephant social behavior, legal elephant hunting, and ecotourism. We encourage the government of Botswana to take immediate action against poaching, including improving monitoring and strengthening transboundary cooperation.

## Introduction

Illegal hunting is a leading threat to wildlife populations worldwide (Ripple et al. 2016; Rija et al. 2020; Paudel et al. 2025). Among land mammals, 17% of endangered species are primarily threatened by poaching (Schipper et al. 2008). Illegal hunting can have a variety of motivations including food acquisition, the bushmeat trade, trade in decorative or medicinal animal products, and human-wildlife conflict (Muth and Bowe Jr. 1998; Janssen et al. 2024). Funds from sale of poached animal products can go to fund regional militias and terrorism (Wang and Tian 2024; Lupton 2025). Thus, combating poaching has important implications for both human and animal populations.

African savanna elephants (*Loxodonta africana*) have been heavily affected by poaching for the ivory in their tusks. Before European colonization, Africa may have had 20 million elephants (Milner-Gulland and Beddington 1993). But unrestrained ivory hunting in the 19^th^ and early 20^th^ centuries and bouts of heavy poaching in the 1970s-1980s and 2000s reduced the population of savanna elephants to ∼350,000 by 2015 (Milner-Gulland and Beddington 1993; Chase et al. 2016). Rapid elephant losses during the 2000s and 2010s finally prompted many countries to ban the ivory trade, and enforcement of anti-poaching laws in Africa has increased recently as well (Zhou et al. 2018; Ngorima et al. 2022). Gains, however, have been uneven (Schlossberg et al. 2020).

Botswana holds Africa’s largest elephant population, ∼140,000 individuals, with ∼95% found in the northern part of the country (Bussière and Potgieter 2023a; Schlossberg and Chase 2024). During 2005-2015, while much of the continent experienced heavy poaching, little poaching appeared to occur in Botswana, and elephant numbers in the country were stable (Chase 2011; Chase et al. 2015). But during a 2018 aerial survey of northern Botswana, we estimated that 385 elephants had been poached for their ivory in the year prior to the survey (Chase and Schlossberg 2019). The poaching appeared to continue into 2019, as more freshly killed elephants were found after the 2018 survey (Schlossberg et al. 2019). These initial reports were publicly disputed and described as "baseless and untrue" by Botswana’s president (Mmolai 2018), making continued, verifiable monitoring of poaching important.

Evidence since 2018-2019 suggests an ongoing poaching problem in Botswana. On aerial surveys of northern Botswana, carcass ratios, the ratio of dead to live-plus-dead elephants, increased steadily from 2010 through 2022, suggesting rising mortality (Schlossberg and Chase 2024). We were not able to conduct surveys for poaching during 2020-2022 when our research permits were suspended by the government of Botswana. When surveillance resumed in 2023, we again found fresh carcasses with missing tusks and extensive cranial or facial mutilation consistent with ivory extraction. Over the subsequent three years, we have located continued evidence of poaching which we report here.

Monitoring elephant poaching in Botswana is difficult because of the size and remoteness of the elephant range. Elephants occupy roughly 100,000 km^2^ in northern Botswana (Thouless et al. 2016), much of which is roadless de facto wilderness (Ibisch et al. 2016). Combating elephant poaching could be easier if anti-poaching forces have a better idea of where poaching is occurring. Past research has shown that poachers often follow predictable patterns in terms of how and where they kill animals (Shaffer and Bishop 2016; Ferreguetti et al. 2018; Kuiper et al. 2020). For instance, Rashidi et al. (2016) found that poachers were most active in areas with higher livestock and waterhole density. Performing a similar analysis of human and environmental factors influencing poaching locations in Botswana could be valuable.

Botswana’s large elephant population and central location within the the five-country Kavango-Zambezi Transfrontier Conservation Area (KAZA), a network of protected areas, make assessing the population’s status critical. Across Africa, poaching has decimated formerly large elephant populations (Chase et al. 2016). Thus, the size of Botswana’s elephant population does not make it immune from poaching. To aid in the fight against poaching, here we provide updated information on locations, timing, and frequency of elephant poaching in Botswana. We also use novel point process models to determine what environmental and anthropogenic factors are associated with poaching.

## Methods

### Study area

Our study area was defined by locations where we located poached elephant carcasses in northern Botswana (Fig. 1). This area was primarily in the Ngamiland district of northern Botswana, including areas north of the Okavango Delta and east of the Okavango Panhandle, extending east to the western portion of Chobe National Park (NP). A small number of carcasses found to the west of the primary mass of carcasses were not included in our spatial analyses, as discussed below.

**Fig. 1.**
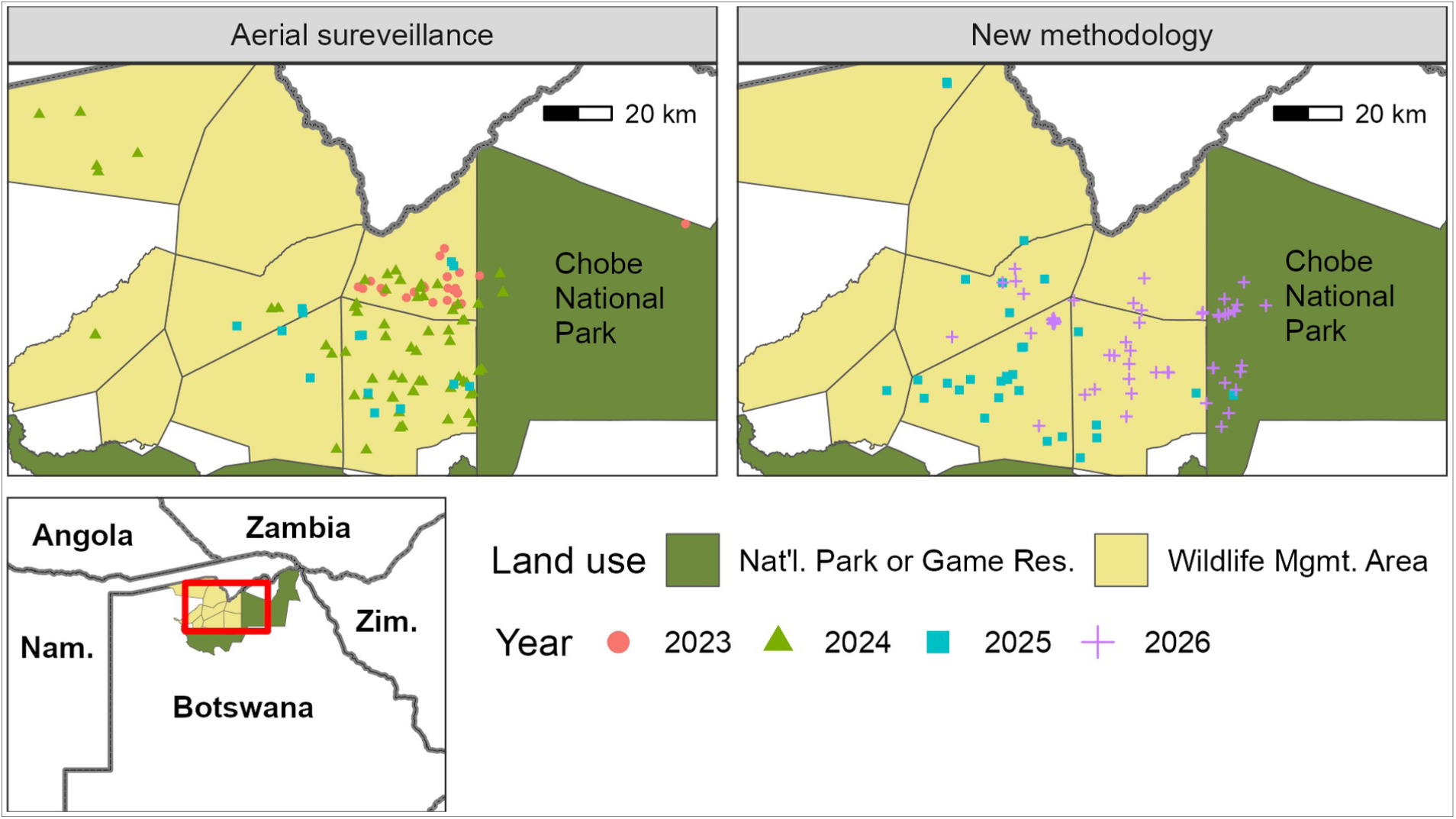
Study area and locations of poached elephant carcasses in northern Botswana, 2023-2026, separated by year and method for locating carcasses. See lower left for context.

The study area is almost entirely undeveloped and includes a variety of habitats including woodlands, shrublands, grasslands, savannas, and marshes. Water is available in a few perennial rivers and, during the wet season, in seasonal pans throughout the area. The area has a subtropical climate with rainfall averaging 460 mm year^-1^ and primarily occurring during a November-April wet season. Per a 2022 aerial survey, the dry-season population of elephants in the study area is approximately 20,000 (Bussière and Potgieter 2023b). Elephant densities in this area are among the highest recorded in Botswana on that survey (Bussière and Potgieter 2023a). The portion of the study area outside of Chobe NP is designated as wildlife management areas, used primarily for photographic safaris. Elephant hunting is not permitted anywhere in the study area. A small number of villages and facilities for tourists are scattered around the study area.

### Locating poached elephants

We used two methods to locate poached elephant carcasses. First, from October 2023 through May 2025, we used intermittent aerial search flights to find poached elephants. Flights were generally conducted in areas where we had previously located poached elephants or where reports from safari guides or law enforcement officials indicated that poaching may have occurred. We searched for carcasses with a Cessna 206 airplane or a Robinson R44 helicopter flown at 60-120 m altitude. When we saw an elephant carcass, we circled over the carcass at a lower altitude to take photographs and obtain geographic coordinates with a GPS receiver. In a few instances, we used the helicopter to visit carcasses on the ground. Carcasses were considered poached if the tusks were missing and the skull or face showed extensive damage consistent with deliberate tusk extraction. Most poached carcasses also were covered with vegetation to conceal them. We classified carcasses as "fresh" or "recent" based on criteria established by CITES’ Monitoring the Illegal Killing of Elephants (MIKE) programme (CITES MIKE Programme 2020). Fresh carcasses had still had flesh and skin, and the ground was still moist from body fluids. These carcasses were likely <1 month since death. Recent carcasses were more decomposed, with little or no flesh remaining, but still had a rot patch around the carcass and skin present.

For November 2025 through July 2026, we used a second detection method to locate poached carcasses. To prevent poachers from circumventing this new method, we cannot include the details of the methods in the main text of this paper. We describe these methods in an appendix which was available to reviewers and is available by request from the authors. We believe that this new method allows us to locate a large majority of poached elephant carcasses in our study area. Once a carcass had been located using the new method, we visited it via aircraft to verify that the carcass was killed illegally, as above.

### Poaching summary statistics

We tallied the number of poached elephants located by year and method. Because detection of poached elephants via aerial surveillance was surely incomplete, observed numbers from that time period should be considered a minimum number poached and not an estimate of the total. We believe that the method used in 2025-2026 has a higher probability of detecting poached carcasses than aerial surveillance, though the exact detection probability is unknown. Regardless, numbers presented in the results are simply observed totals and not estimates of total numbers poached. To determine how locations of poaching changed from those observed in 2018, we mapped 2023-2026 poaching locations against 2018 hotspots from Schlossberg et al. (2019).

Botswana’s climate is highly seasonal, and little is known about when poachers are most active within the year. Thus, we were interested in assessing the seasonality of poaching. Unfortunately, neither of our sampling methods were well-suited for statistical analysis of poaching dates. Aerial surveillance was intermittent and took place on only 16 dates over 2 years. The newer method was continuous and should be more likely to detect poached elephants but was only used from November 2025 through 1 July 2026, which was the cutoff date for inclusion in this study. Consequently, we did not have a full year of data from the new method to analyze the seasonality of poaching. Instead, to illustrate when we located carcasses, we plotted the number of "fresh" carcasses located via each method by month. Fresh carcasses were likely dead for less than a month at the time they were located, so the date they were found should be a good approximation of the poaching date for assessing seasonality.

### Spatial models for predicting poaching locations

To aid in anti-poaching efforts, we used spatial models to identify predictors of poached carcass density and to map predicted carcass density in northern Botswana. Given that elephants are widespread in our study area, carcass density should be an index of poaching risk. We used Poisson point process models (PPMs) to model the intensity of elephant poaching. PPMs model point data as the realization of a Poisson process with a spatially varying mean known as intensity (Baddeley et al. 2015). Intensity is the expected density of poached carcasses.

Previously, point process models have been used to examine clustering in poaching locations. To our knowledge, ours is the first study to use point process models to predict the intensity of poaching as a function of spatial covariates. As discussed below, PPMs are flexible and relatively easy to implement, making them well-suited for analyzing poaching.

Ignoring constants, the log likelihood for the PPM is

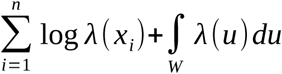

where *λ* is the estimated intensity at the location specified in parentheses, *x_i_* are carcass locations, *n* is the number of carcasses, *W* is the study area, and *u* is a point in *W* (Baddeley et al. 2015). The integral in the likelihood is intractable but can be estimated with numerical quadrature, measuring *λ* at a large number of random or systematic background points (Renner et al. 2015). For our models, we placed quadrature points on a grid with 200 meter spacing.

Intensity in a PPM can be modeled as a log-linear function of parameters so that

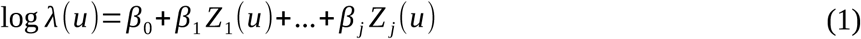

where *j* is the number of parameters, *Z* (*u*) denotes a covariate measured at location *u*, and *β*_0_ … *β _j_* are parameters to be estimated, including an intercept. With a sufficiently large number of quadrature points and appropriate weighting of observations, the likelihood of the PPM becomes equivalent to that of a Poisson generalized linear model (GLM; Berman and Turner 1992).

Accordingly, the PPM can be modeled as

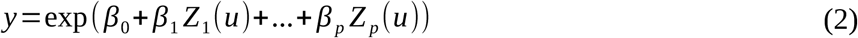

where *y* is 0 for quadrature points and 10^6^ for carcass locations (Dovers et al. 2024). The resulting log likelihood is

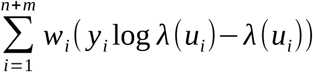

where *m* is the number of quadrature points and *w*= 1/[area of study area] for quadrature points or 10^-6^ for carcass locations.

PPMs generally assume that intensity is independent after conditioning on predictors (Baddeley et al. 2015). With elephant poaching, independence is likely to be violated because multiple elephants may be killed by the same poachers acting locally. In fact, we located 33 pairs of carcasses <1 km from one another that were likely killed no more than a few days apart.

Exploratory analysis with inhomogeneous PPMs revealed significant clustering of carcasses at distances of 0-7,500 m. To account for this lack of independence, we modeled the point process for poached elephants as a log-Gaussian Cox process (LGCP). LGCPs are the equivalent of a linear model with a random intercept, allowing background intensity to vary spatially via a random Gaussian process (Baddeley et al. 2015). Accordingly, the right side of Eq. 2 was modified to add a spatially correlated random effect with mean of 0. The covariance of the random effect decreases with distance between points. The result is a background intensity that is "clumpy" and more likely to produce clustered observations than a PPM with a constant intercept.

Based on past research on elephant poaching and our experience in Botswana, we identified 12 variables that potentially predicted poaching intensity (Table 1; see Appendix S1 for details). We included tree cover and EVI, a measure of vegetation greenness and leaf cover, because poachers might seek out areas with more vegetation to avoid being detected or to match elephants’ habitat preferences (Maingi et al. 2012). Human population and distance to road measured accessibility by local inhabitants or people traveling in vehicles (Shaffer and Bishop 2016). Distance to BDF patrol bases could be important if poachers are avoiding anti-poaching units (Leader-Williams et al. 1990). We included distance to the Namibian border because authorities in Botswana and Namibia have intercepted poachers crossing from Botswana into Namibia with elephant tusks. Distance to water was included because we have found several poached elephants near waterholes, possibly indicating that poachers are waiting there for bulls to visit (Kuiper et al. 2020). Slope and elevation were included to test whether topography influenced poaching locations (Beale et al. 2018). Maps of predictor variables are available in Fig. S1 in Appendix S2.

**Table 1.** Variables included in spatial models of elephant poaching in Botswana. See Appendix 1 for details.

| Variable | Description |
|---|---|
| Distance to road | Distance to nearest road |
| Distance to border | Distance to Namibia border |
| EVI (wet) | Wet-season Enhanced Vegetation Index |
| EVI (dry) | Dry-season Enhanced Vegetation Index |
| Trees | Tree cover as a % of land area |
| Humans (1 km) | Est. human population within 1 km of point |
| Humans (10 km) | Est. human population within 10 km of point |
| Distance to water (wet) | Distance to surface water, wet season |
| Distance to water (dry) | Distance to surface water, dry season |
| Distance to BDF camp | Distance to Botswana Defence Force patrol camp |
| Slope | % slope of terrain |
| Elevation | Elevation in m |

Because we used two distinct methods to locate carcasses, the possibility exists that the variables above could have different effects on carcass density depending on the method used to find carcasses. Thus, we ran three sets of models, one for carcasses located via aerial surveillance, one for carcasses located using our new method, and one for all carcasses combined.

For each of the three datasets, we used the following procedures. First, we identified the study area as the minimum convex polygon containing the center-most 90% of carcass locations. We buffered this polygon by 1 km to avoid edge effects in the fitting of the random effect. To identify the selected carcasses, we calculated the distance from each carcass to the centroid of all carcass locations and took the lowest 90% of distances. We did this because we located a few outlying carcasses to the west of the main study area (Fig. 1). Including such carcasses would have expanded the study area manyfold and resulted in large areas with no carcasses. We were interested in understanding how poachers are operating within a poaching hotspot. If we had including outlying carcasses, the model might have primarily served to distinguish areas with poaching from the larger area where poaching was not occurring. Additionally, the outlying areas with carcasses had much lower elephant densities than the primary study area (Bussière and Potgieter 2023b). Accurately modeling poached elephant density in the larger area would have required controlling for elephant density, and the latest survey data available were from 2022 and may no longer be accurate.

To fit models, we took advantage of the fact that the Gaussian random field for the LGCP can be estimated using standard software for fitting generalized additive models (Dovers et al. 2024). Accordingly, we used the *mgcv* software package to fit models (Wood 2017). We modeled the random field as a two-dimensional Gaussian process smooth as suggested by Dovers et al. (2024). We set the number of knots in each dimension to 300 to ensure that the number of knots was much larger than the estimated degrees of freedom of the final smooth.

For the predictors in Table 1, we took advantage of the fact that in Eq. 1, the terms for individual variables can also be non-linear functions. Thus, to allow for non-linear relationships between predictors and intensity, we modeled each variable as a cubic regression spline with three knots. This allowed smooths to be linear or have a simple curved shape. To aid in variable selection, we used the shrinkage version of the regression splines from *mgcv*. This formulation shrinks smooth components towards zero and is roughly equivalent to a LASSO penalty (Marra and Wood 2011). This penalization allowed us to combine parameter estimation and variable selection in a single model run. In general, using shrinkage splines in our models resulted in estimated smooth terms being linear rather than curved. We fit the Gaussian process smooth without shrinkage because we used a separate procedure to prevent overfitting the random effect, as below.

Within the study area, we generated quadrature points on a 200-m grid. We measured all spatial covariates in Table 1 at each carcass location and quadrature point. To prevent correlated predictors from influencing results, we first ran a basic inhomogeneous point process model with all variables included but no random spatial effect. We calculated each variable’s concurvity, a non-linear equivalent of multicolinearity. We sequentially removed the predictor with the highest concurvity until no predictors with concurvity >0.5 remained. Remaining predictors were used for in the final models described below.

The remaining parameter to be estimated in our models was *ρ*, the range parameter defining the covariance of the Gaussian random field. Essentially, *ρ* defines the "roughness" of the Gaussian field; larger *ρ* values produce smoother surfaces. As a general rule, likelihood-based criteria tend to favor rougher Gaussian fields with smaller *ρ*, but such fields are likely to be overfit and have poor predictive ability. To determine the optimal *ρ* for our models, we ran models across a wide range of *ρ* values and compared the generalized cross-validation score across models (Dovers et al. 2024). This score penalizes roughness in the Gaussian field, leading to a better balance between model fit and predictive ability (Wood 2017). To find the optimal *ρ* value for each dataset, we ran models with *ρ* values of 100 m as well as 1,000 m through 10,000 m in increments of 1,000 m and 20,000 m through 100,000 m in increments of 10,000 m. We also tested a model with no pre-set value of *ρ* in which case *mgcv* defaults to using the largest spacing between points, which was 68-85 km in our models. For all three models, GCV scores were clearly lowest for *ρ* < 3000 m. We ran with models *ρ* = (100, 200, …, 3000) and used the single model with the lowest GCV score for inference.

We used two methods to validate each final model. First, we calculated the inhomogeneous K function for the final model and compared the observed value against values from simulated point processes based on the predicted intensity at the quadrature points. The K function measures the clustering or dispersion of a point process relative to its intensity (Baddeley et al. 2015). Using the *spatstat* package (Baddeley and Turner 2005), we simulated 1,000 point datasets from the predicted intensity map and calculated K as a function of distance for each dataset. Observed K values outside of the 95% confidence intervals of the simulated values indicate clustering (for higher K values) or dispersion (for lower K values) of points beyond what the model predicts.

Second, to assess model fit, we used 5-fold cross validation (Boyce et al. 2002). We randomly divided the dataset into five folds while ensuring that each fold contained the same number of quadrature points and the same number of carcasses, to ± 1 per group. We serially excluded each fold from the dataset and then ran the final model as above. We then used the models to predict *λ* for the excluded points. We combined the predicted values from the five folds, divided that dataset into 20 bins by predicted *λ* value, and calculated *p*, the proportion of points that were carcasses, for each bin. We used Spearman’s *ρ* to calculate the rank correlation between bin number and *p*. If the model fits the data well, higher deciles of *λ* should have more carcasses, resulting in a positive correlation. We considered the model to fit well if Spearman’s *ρ* was significantly greater than 0.

## Results

From October 2023 through early July of 2026, we located 202 poached elephants (Table 2). Of this total, we found 118 via aerial surveillance, and 84 via the new method. As noted above, the new method was only in place from November 2025 through 1 July 2026. Compared to 2018 poaching hotspots, heavily overlapped the NG15 hotspot from 2018 (Fig. 2, central hotspot). The area between that hotspot and the NG11 hotspot to the west had no poaching in 2018 but numerous carcasses in 2023-2026. The eastern hotspot in Fig. 2 had no poached carcasses in the current study, though one carcass was located just outside of it.

**Fig. 2.**
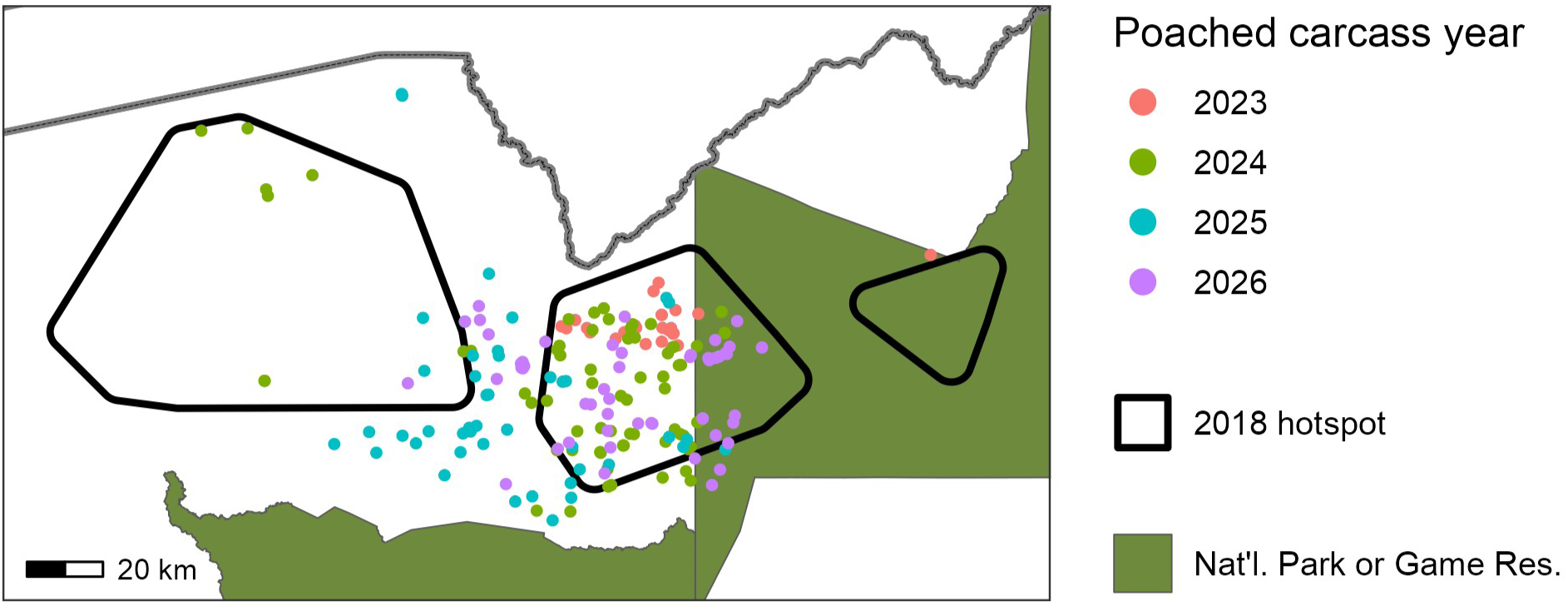
Comparison of 2018 poaching hotspots and 2023-2026 poached carcass locations. See Fig. 1 for context.

**Table 2.**
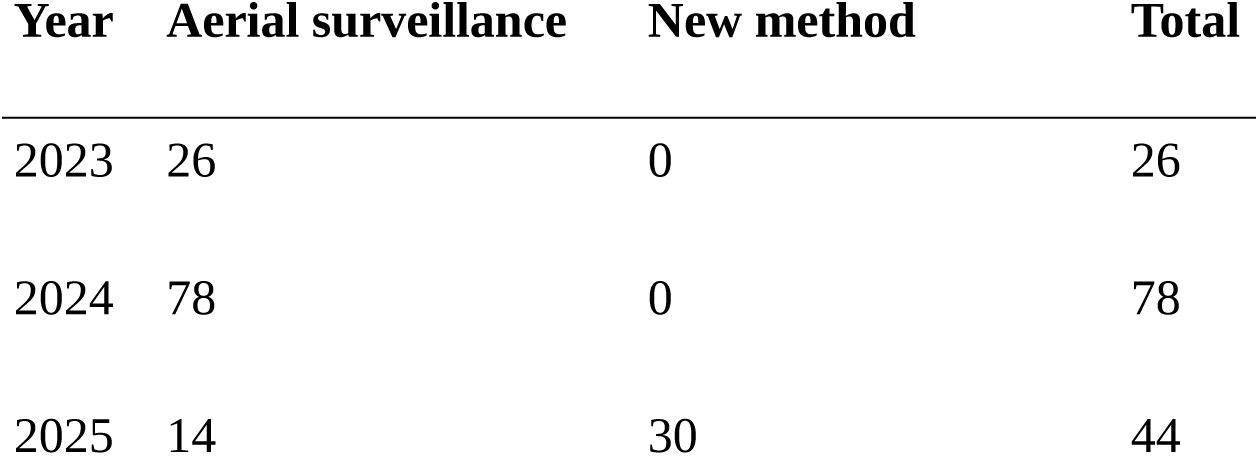
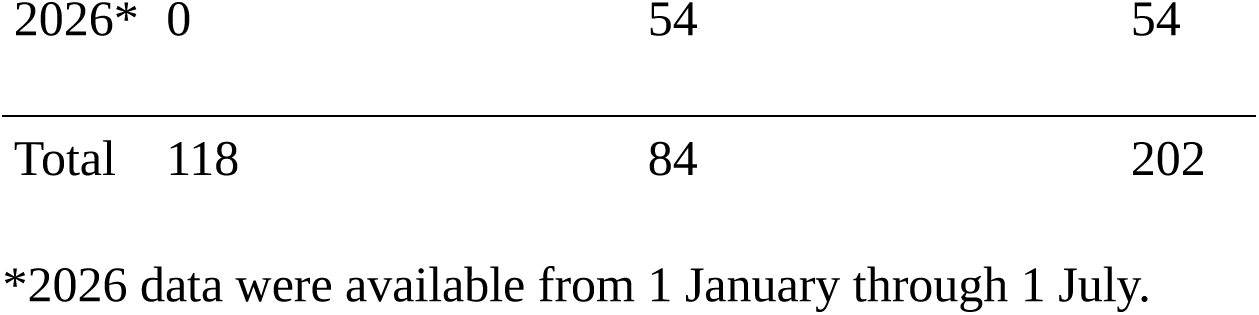
Number of illegally killed elephants located in northern Botswana by method and year of location.

| **Year** | **Aerial surveillance** | **New method** | **Total** |
|---|---|---|---|
| 2023 | 26 | 0 | 26 |
| 2024 | 78 | 0 | 78 |
| 2025 | 14 | 30 | 44 |
| 2026* | 0 | 54 | 54 |
| Total | 118 | 84 | 202 |

\*2026 data were available from 1 January through 1 July.

For fresh carcasses, likely killed within a month of being located, we did not observe any obvious seasonality, though no statistical test was possible due to restricted sampling dates (Fig. 3). Based on the sizes of the carcasses and the exposed penis visible in many carcasses (which typically occurs when elephants experience pain), all carcasses poached were classified as adult males.

**Fig. 3.**
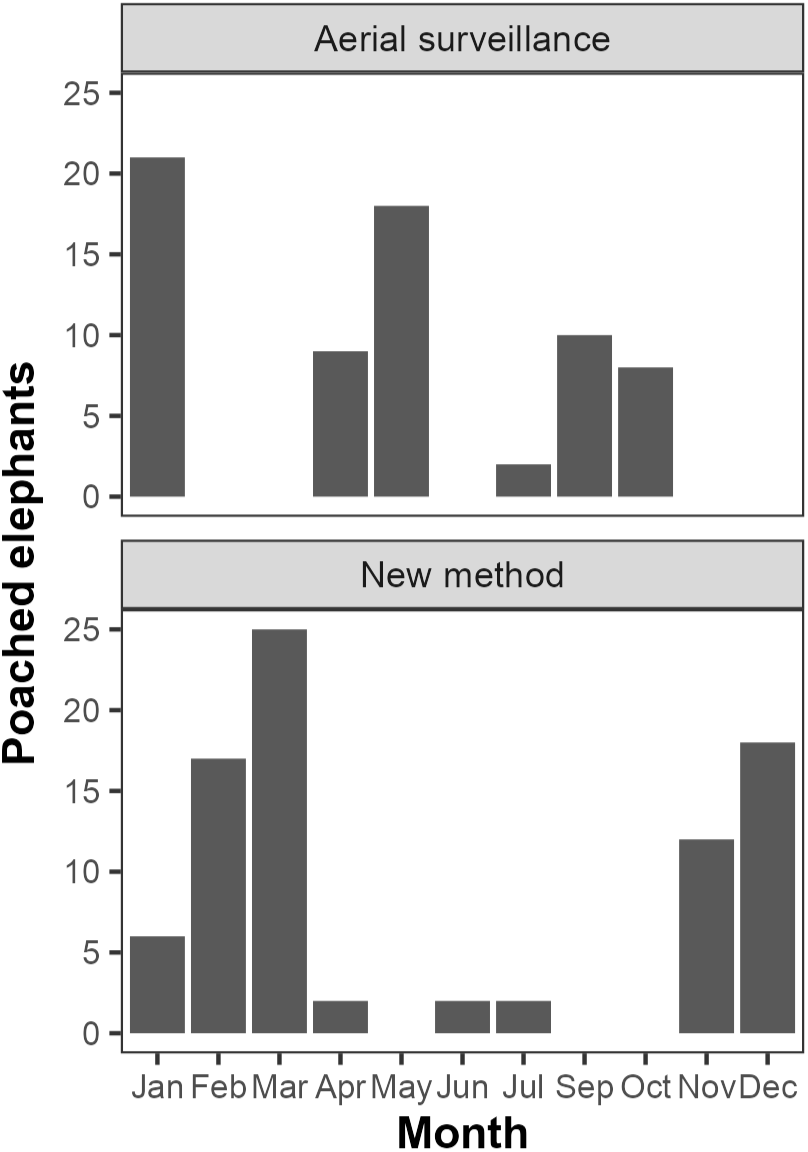
Number of fresh carcasses located by month, northern Botswana, 2023-2026.

We used PPMs to model factors influencing carcass locations for three datasets: 1) carcasses located via aerial surveillance, 2023-2025 (*n* = 96; study area size = 2,647 km^2^); 2) carcasses located via the new method, 2025-2026 (*n* = 79; 4,227 km^2^); 3) all poached carcasses in our dataset, 2023-2026 (*n* = 186; 4,159 km^2^). Outlying carcasses were not included in these datasets, so numbers of carcasses are lower than the numbers in Table 2. The optimized range parameter, *ρ*, for the LGCP was 400 m for aerial surveillance and 1,200 m for the other two datasets.

The two methods for locating carcasses produced distinct results (Table 3). For aerial surveillance, carcass density decreased with distance to BDF base and showed a parabolic effect of distance to road with density highest at distances of ∼7 km (Fig. 4). For carcasses located via the new method, density increased with slope and distance to the Namibia border (Fig. 5). For all carcasses combined, slope and distance to road had effects similar to those from the first and second models above, respectively (Fig. 6). In this model, carcass density also increased with dry-season EVI.

**Fig. 4.**
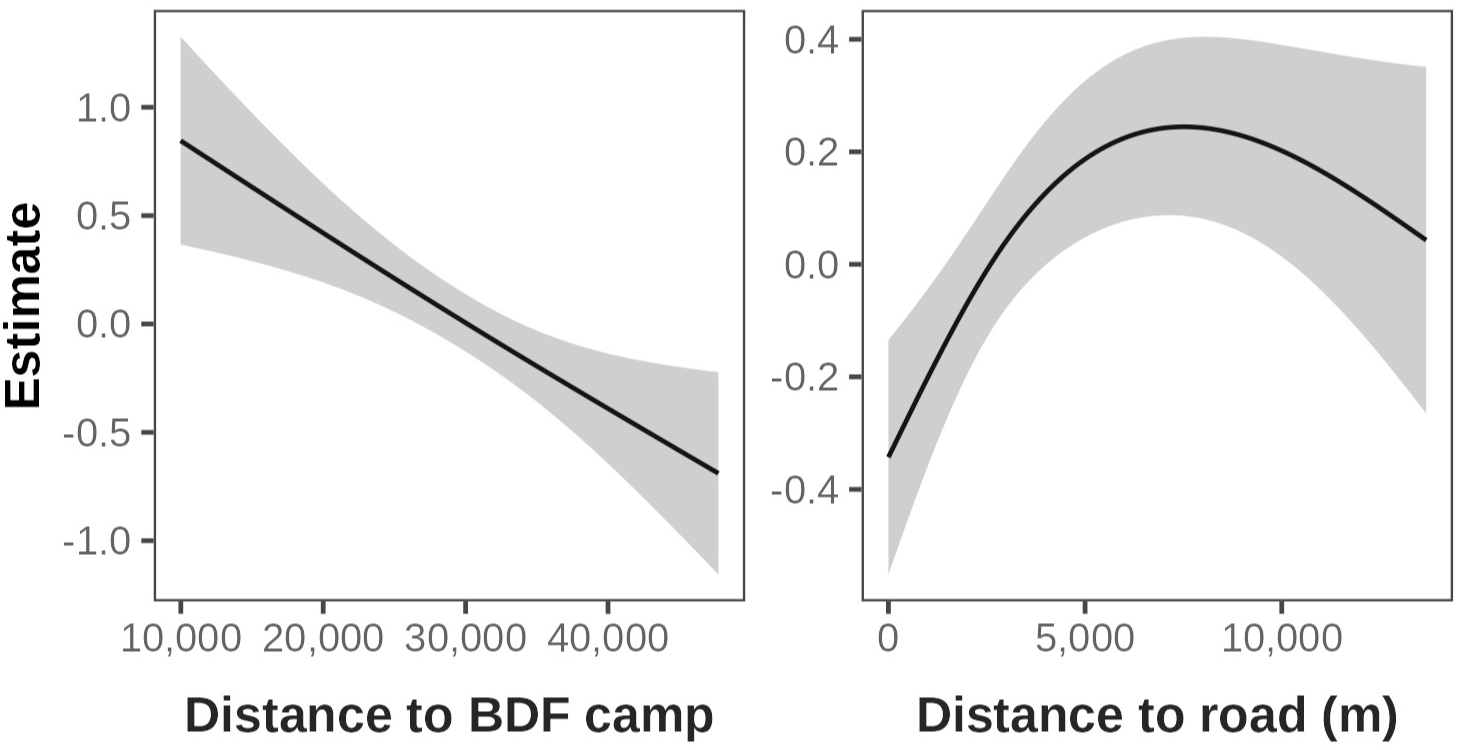
Predicted effects for significant terms in point process models predicting poached carcass density for carcasses found via aerial surveillance, Botswana, 2023-2025.

**Fig. 5.**
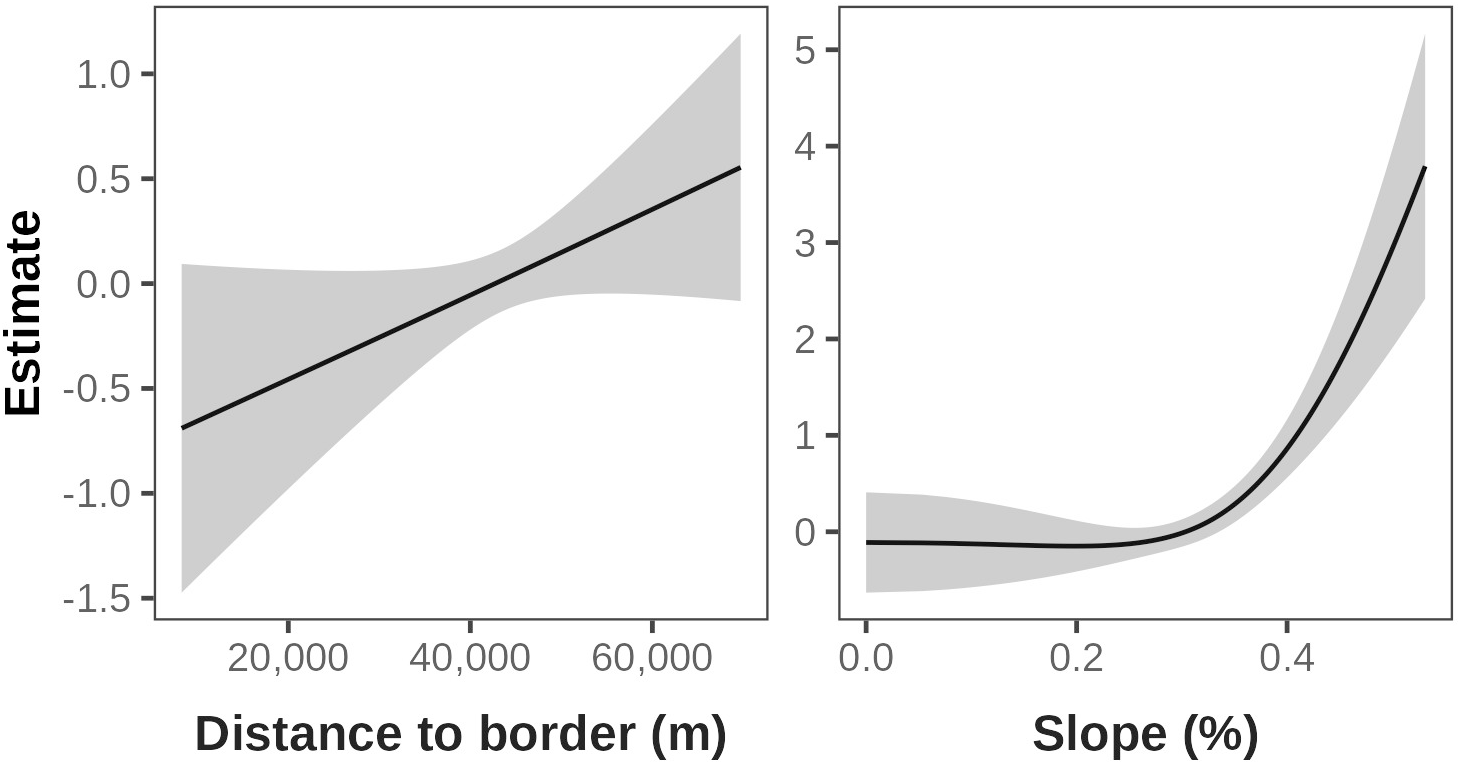
Predicted effects for significant terms in point process models predicting poached carcass density for carcasses found via the new method, Botswana, 2025-2026.

**Fig. 6.**
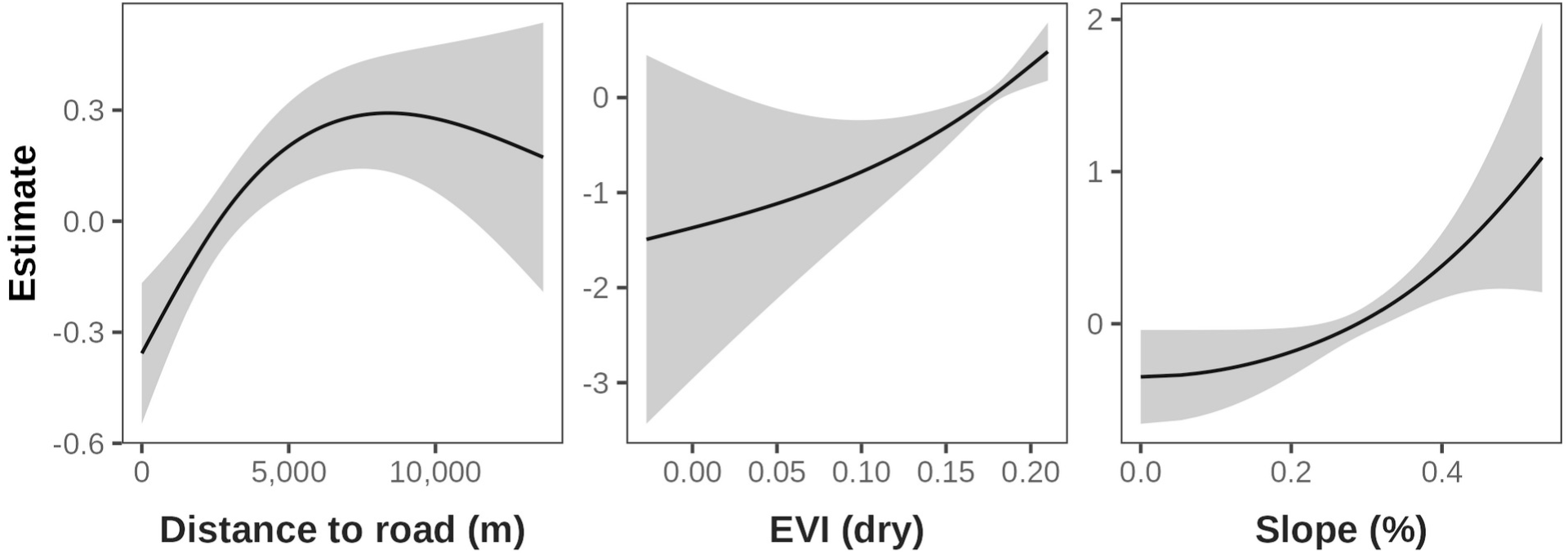
Predicted effects for significant terms in point process models predicting poached carcass density for all carcasses, Botswana, 2023-2026.

**Table 3.** Results from point process LGCP models predicting poached carcass intensity as a function of spatial predictors. "Method" column indicates the method used to locate carcasses.

| **Method** | **Parameter** | **Est. df** | **χ<sup>2</sup>** | **P** |
|---|---|---|---|---|
| Aerial surveillance | **Distance to road** | **0.61** | **4.53** | **0.001** |
|  | Distance to border | 0.25 | 0.46 | 0.09 |
|  | EVI (wet) | 0.92 | 1.37 | 0.17 |
|  | EVI (dry) | 0.94 | 0.91 | 0.30 |
|  | Trees | 0.64 | 0.74 | 0.27 |
|  | Humans (1 km) | 0.52 | 0.66 | 0.24 |
|  | Distance to water (dry) | 0.50 | 1.40 | 0.05 |
|  | **Distance to BDF camp** | **1.20** | **3.86** | **0.02** |
|  | Elevation | 0.54 | 1.23 | 0.07 |
|  | Slope | 0.90 | 0.44 | 0.56 |
|  | LGCP random effect | 52.43 | 80.52 | 0.76 |
| New method | Distance to road | 0.45 | 0.56 | 0.13 |
|  | **Distance to border** | **0.45** | **0.84** | **0.03** |
|  | EVI (wet) | 0.44 | 0.16 | 0.50 |
|  | EVI (dry) | 0.46 | 0.96 | 0.10 |
|  | Trees | 0.15 | 0.02 | 0.69 |
|  | Humans (1 km) | 0.31 | 0.10 | 0.55 |
|  | Distance to water (dry) | 0.35 | 0.35 | 0.22 |
|  | Elevation | 0.12 | 0.06 | 0.50 |
|  | **Slope** | **1.97** | **9.44** | **0.01** |
|  | LGCP random effect | 81.82 | 140.50 | 0.27 |
| All methods | **Distance to road** | **0.90** | **5.25** | **0.0004** |
|  | Distance to border | 0.09 | 0.04 | 0.35 |
|  | EVI (wet) | 0.78 | 1.92 | 0.07 |
|  | **EVI (dry)** | **0.82** | **2.84** | **0.03** |
|  | Trees | 0.07 | 0.02 | 0.55 |
|  | Humans (1 km) | 0.09 | 0.06 | 0.41 |
|  | Distance to water (dry) | 0.22 | 0.30 | 0.14 |
|  | Elevation | 0.21 | 0.09 | 0.47 |
|  | **Slope** | **1.12** | **3.68** | **0.05** |
|  | LGCP random effect | 86.14 | 117.62 | 0.90 |

For carcasses detected via aerial surveillance, predicted intensity was generally highest in the southernmost part of the study area (Fig. 7a). For carcasses located with the new method, predicted intensity was highest in the southern and southwestern part of the study area (Fig. 7b). When we combined all carcasses, the effects of roads were paramount, as indicated by the darker corridors in Fig. 6c (see Fig. S1). For the latter two models, the effects of slope were illustrated by the line of predicted high carcass density in the eastern part of the study area, an area of high slope relative to the remainder of the study area (Fig. S1).

**Fig. 7.**
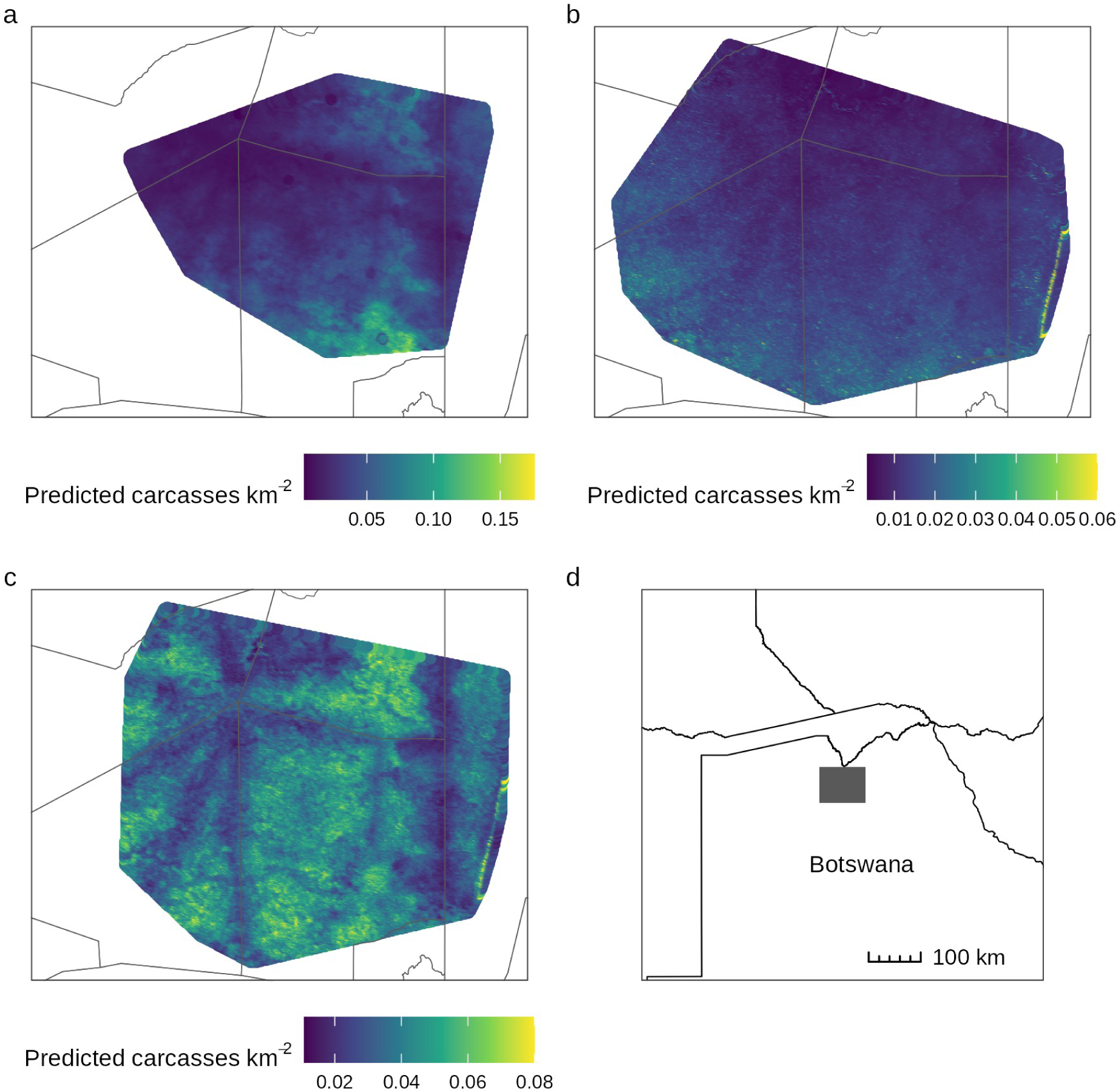
Maps of predicted poaching intensity for carcasses detected via (a) aerial surveillance, (b) the new method, and (c) both methods combined. These plots show predictions from fixed effects alone; random effects were excluded. Shaded area in (d) is the plotted area for (a), (b), and (c). Thin lines are boundaries of Controlled Hunting Areas in Botswana.

All three models showed a good fit to the data per five-fold cross validation (all ρ ≥ 0.56, all P < 0.01). K-function plots showed that the dispersion of carcasses was consistent with predicted intensity for all three models (Fig. S2 in Appendix S2). Interestingly, all three models showed a slight clustering of carcasses, beyond what the models predicted, at a distance of ∼2-3 km. This could indicate the scale at which poachers were operating (see Discussion).

## Discussion

The 202 poached elephants recorded demonstrate that illegal killing of elephants for ivory has continued in northern Botswana after the 2017-2019 outbreak. Based on estimated carcass sizes and ground examination of some carcasses, all poached animals in this study were classified as adult males. This is consistent with the earlier outbreak in which all verified carcasses were mature males (Schlossberg et al. 2019). Selective killing of bulls suggests that poachers are targeting them for the ivory in their relatively large tusks (Whyte and Hall-Martin 2018). Elsewhere in Africa, poachers have been less selective and have also killed family groups comprised mainly of females and young (Wittemyer et al. 2013; Jones et al. 2018). For some populations, this non-selective poaching led to severe losses (Chase et al. 2016). No such broad-scale poaching has been observed in Botswana to date; poachers appear to be continuing to hunt adult males.

Based on a 2018 aerial survey, we estimated that 385 elephants had been poached in Botswana in a roughly one-year period (Schlossberg et al. 2019). Because the current study did not use systematic sampling, and detection probabilities for poached carcasses were unknown, we cannot estimate total numbers poached for 2023-2026. But the newer method used in 2025 and 2026 resulted in 96 carcasses being located in 7.5 months. If this rate were to continue for a full year, this would result in 154 poached carcasses being located. Again, however, these data are not sufficient to compare the current poaching rate with the 2018 sample.

Compared to 2018, poaching in 2023-2026 appears to have shifted in location (Fig. 2). The NG15 hotspot had extensive poaching in both 2018 and the current study. But the other two hotspots had only a few carcasses in the current study. Thus, poaching appears to have changed locations during the five years we were unable to monitor carcasses. Reasons for this shift are unknown, but the PPMs also showed that poacher behavior in Botswana may have changed over time (see below).

Because the government of Botswana rescinded our research permit, we were unable to monitor poaching from 2020 through late 2023. We believe that the most parsimonious explanation for the poaching observed in this study is that the poaching we documented in 2018 and 2019 continued through the period when no monitoring was occurring. The increase in elephant carcass ratios in Botswana between 2018 and 2022 is consistent with this hypothesis (Schlossberg and Chase 2024).

### Point process models of poached carcass intensity

Each of three datasets led to a different predicted pattern of poached carcass intensity (Fig. 7). For carcasses detected with serial surveillance, the highest predicted densities were in the east-central and southernmost portions of the study area. The new method had the highest modeled intensities in the southern and southwestern part of its study area. When we combined all carcasses, model predictions did not show any obvious geographic gradient, though avoidance of roads was apparent.

The three models differed in the variables that affected carcass density. For the carcasses detected aerially, distance to BDF camps and distance to roads were the only significant predictors. We expected that poachers might avoid BDF camps, but carcass density actually increased nearer camps. No BDF camps were located within our study areas, and observed distances to camps were 10-40 km for carcasses (Fig. S1). If poachers were avoiding patrol bases, it might have been scales smaller than 10-40 km. Interestingly, Maingi et al. (2012) also found that poaching rates were higher closer to ranger bases in Kenya, though they did not report the scale of this effect.

We found significant effects of distance to road in the aerial surveillance model and for all carcasses combined. For both models, carcasses were less common within ∼5 km of roads. Roads are uncommon in our study area (Fig. S1), so poachers may be traveling on foot and staying away from roads to avoid detection. Past studies have found inconsistent effects of roads on elephant poaching. Maingi et al. (2012) found a positive correlation between road density and poaching rates in Kenya. For a different time period, Rashidi et al. (2016) found no significant effects of roads on poaching in the same ecosystem.

Carcass density increased with slope for carcasses detected via the new method and for all carcasses combined. Only one small portion of the study area, at the far eastern edge, had slopes >0.4%; the remainder of the study area was nearly flat (Fig. S1). Several poached carcasses located via the new method were found in the area with relatively high slopes. Because only one portion of our study areas had the higher slopes, determining whether this result reflects tactics used by poachers or was spurious will require additional study.

For carcasses located via the new method, density increased with distance from the Namibia border, creating a rough north-south gradient in predicted intensities (Fig. 7b). And for all carcasses combined, carcass density increased with dry-season EVI. This could reflect poachers seeking area with greater vegetation cover during the dry season when many trees and shrubs in the study area leafless.

We found no significant effects of distance to water, tree cover, elevation, or human population in our study. Some of these variables did predict poaching in other studies. For instance, Beale et al. (2018) found that poaching was associated with lower elevation and intermediate distance to water. And Rashidi et al. (2016) found more poaching in areas with higher waterhole density. This, combined with the fact that our three models differed in both their spatial predictions and the variables that affected carcass density, suggests that the factors affecting poaching locations are something of a moving target. One possible explanation is that poacher tactics change over time or depending on the group doing the poaching. Beale et al. (2018) found some differences in predictors of poaching between years in Tanzania. Maingi et al. (2012) identified different predictors of poaching between wet and dry seasons. Changes in poacher behavior over time or due to seasonal factors would make it more difficult to predict poaching intensity and combat poaching.

Kirkland et al. (2025) used simulations to show that poaching could be modeled as a point process. To our knowledge, however, ours is the first field study to predict poaching locations as a function of covariates using point process models. PPMs proved well-suited for analyzing such data, with models showing a good fit and identifying several important covariates. The methods described here are flexible and relatively easy to implement while accounting for the spatial autocorrelation between carcass locations (Dovers et al. 2024). Thus, we suggest that future analyses of poaching incorporate PPMs if possible.

### Poaching tactics

For Botswana, we can make a few conclusions about potential tactics being used by poachers. First, we believe that poachers are largely operating on foot when shooting elephants. Median distance to road for poached carcasses in our dataset was 3.8 km. We have not observed vehicle tracks near any of the poached carcasses that we located. During aerial surveillance we have located a few temporary poachers’ camps. Poachers could be establishing temporary camps and then searching for elephants in the surrounding area. This would explain the clustering of carcasses on scales of a 2-3 km that was not fully explained in our models (Fig. S2).

### Impacts of poaching

The total of 202 poached elephants from this study is just 0.14% of Botswana’s estimated total of 140,000 elephants as of 2022 (Schlossberg & Chase 2024). Because, however, poachers appear to be targeting mature males in a relatively small portion of northern Botswana, impacts may be disproportionate to the numbers taken. Male elephants have lower survival rates than females, so older males tend to be uncommon in elephant populations (Moss 2001; Wittemyer et al. 2013). In two Tanzania populations little affected by poaching or legal hunting, males ≥25 years old made up just 6.2% of the population (Jones et al. 2018). Thus, poaching concentrated on adult males in a relatively small area could lead to their local depletion. This would be problematic because adult male elephants perform several important behavioral roles in savanna elephants including providing a large share of matings, controlling aggression in younger males, and acting as leaders in groups of traveling males (Poole 1989; Slotow et al. 2000; Allen et al. 2020).

Another potential impact of poaching in Botswana is the loss of males with large tusks. These "tuskers" are important draws for tourism and for elephant hunting (Moller et al. 2019), both of which are important economically in Botswana (Ketshabile and Ferreira 2009). Though legal elephant hunting does not occur in our study area, there are hunting blocks ∼25 km to the southeast of the study area (Schlossberg and Chase 2025). Depending on movements of elephants, depletion of mature males in our study area could negatively impact hunting nearby (Schlossberg and Chase in press). Another potential impact of selective poaching of mature males is genetic selection for smaller tusks or tusklessness. Chiyo et al. (2015) found that tusk length decreased by ∼25% after a period of intense poaching in Kenya.

### Reassessment of 2018 poaching

Following the publication of our initial report on elephant poaching (Chase and Schlossberg 2019), a number of researchers and prominent government officials dismissed the findings. The president of Botswana called the claims "baseless" (Mmolai 2018), and three researchers said that a media report based on the findings had "no scientific basis" (Alexander et al. 2018). We subsequently published our findings in a peer-reviewed article that included dozens of photographs of poached elephants and all of the associated data that we collected (Schlossberg et al. 2019). Still, some researchers, despite overwhelming evidence to the contrary, continued to dismiss our findings (Cassidy and Salerno 2020). The current study should put any doubts about poaching in Botswana to rest. The repeated discovery of illegally killed elephants in the same area identified as a poaching "hotspot" in 2018 demonstrates that the original warning was neither fabricated nor transient.

More recent government statements recognize several of these concerns. In 2025, Botswana adopted a National Anti-Poaching Strategy for 2025–2030, emphasizing inter-agency coordination, intelligence gathering, technological capacity, and transboundary cooperation (Government of Botswana 2025). During parliamentary debate in April 2025, government representatives stated that the previous administration had disarmed wildlife officers, that the absence of appropriate weapons had limited their ability to deter and respond to armed poachers, and that poaching incidents were increasing (Galeragwe 2025). In March 2026, however, the Ministry of Environment and Tourism reported that rearming the Anti-Poaching Unit had “stalled due to budgetary constraints” and had been deferred to a future fiscal year (Ministry of Environment and Tourism 2026).

### Recommendations

With elephant poaching continuing in northern Botswana, we recommend that the country implement a national carcass-monitoring and response system. Such a system would combine standardized aerial surveillance, continuous field reporting, CITES MIKE-compatible carcass classification, rapid ground verification, secure sharing of locations, and annual publication of aggregated mortality data. Tourism operators, concessionaires, professional guides, lodge staff, and local communities are often well-placed to detect carcasses, gunshots, or other indicators of illegal activity. An automated reporting mechanism, with defined response times and protection for informants and commercially sensitive information, could increase reporting levels for poaching incidents. Participation in such a system could become a requirement for operators of tourism concessions.

Because the northern Botswana elephant population forms part of the transboundary Kavango–Zambezi landscape (Lindsay et al. 2017), anti-poaching activities cannot end at national borders. Botswana, Angola, Namibia, Zambia and Zimbabwe should cooperate to integrate incident locations, arrest and prosecution records, ballistics, communications and financial investigations, vehicle movements, and forensic examination of recovered ivory. DNA analyses can identify the geographic origin of seized ivory and link tusks from different seizures to common trafficking networks (Wasser et al. 2015). Such analyses would help determine whether poaching incidents presently treated as isolated events involve recurring personnel, routes, financiers or facilitators.

Finally, we hope that Botswana will be transparent about poaching and publish annual performance indicators. These indicators should include verified carcasses, response times, patrol effort and coverage, ivory recoveries, arrests, prosecutions, and convictions. Transparent reporting should protect exact operational locations while allowing citizens, researchers, and conservation partners to assess whether government commitments are producing measurable results. Elephants are the national heritage of Botswana, and the government should share information about elephant poaching both to allow the public to be informed about the status of this key species and to allow people to hold the government accountable for its actions on poaching.

## Conclusions

Africa’s monitored savanna elephant populations declined by approximately 30% between 2005 and 2015, primarily because of ivory poaching (Chase et al. 2016). Botswana retains the continent’s largest national elephant population and is, therefore, central to the future of the species. Yet national abundance should not obscure persistent illegal killing in localized hotspots nor the selective loss of mature, large-tusked bulls. Despite the warning we issued in 2018, its subsequent peer-reviewed verification, and the additional evidence presented here, elephants continue to be killed for ivory with alarming regularity in the same broad area. We hope that the evidence presented here will generate not only greater public awareness but a measurable and sustained government response. Botswana should welcome scientific, technical and financial assistance from qualified researchers, local communities, tourism operators, and international partners. Transparency and cooperation should not be regarded as threats to the country’s reputation: continued under-reporting, delayed action, and preventable wildlife losses present the greater danger. The question is, therefore, no longer whether the problem exists, but whether this accumulated evidence will finally be met with the urgency, openness, and sustained action necessary to stop it.

## Acknowledgments

We thank Prince Harry, The Duke of Sussex, Claire Bracebridge, Mary Rice, Rodger Schlickeisen, Ramalepa and Associates, Samuel Wasser, Tammi Seaman, and Kurt Steiner for their encouragement and support.

## Appendix S1. Supplemental methods

### Enhanced Vegetation Index (EVI)

EVI data were derived from the MODIS sensor on the Terra satellite and were downloaded from https://lpdaac.usgs.gov/dataset_discovery/modis/modis_products_table/mod13a3. We used the 250 x 250 m gridded product available in 16-day increments. Images used were acquired between 1 January 2023 and 30 June 2026. All data passed quality checks for cloud cover and shadows. In our study area, EVI differs reliably between wet seasons, when vegetation is green, and dry seasons, when herbaceous vegetation goes dormant and many woody plants lose their leaves. To account for these differences, we calculated EVI separately for wet and dry seasons. For each grid cell and year, we calculated the minimum (dry-season) and maximum (wet-season) EVI. Wet-season EVI peaked in January or February of each year, so each wet season (November through April) contributed only one value to this analysis. The wet-season EVI used in our analyses was the mean of the maxima across the four years by grid cell. The dry-season EVI was the mean of the minima by grid cell.

### Distance to road

We obtained a shapefile of OpenStreetMaps (OSM) roads in Botswana from https://download.geofabrik.de/africa/botswana.html. OSM data are contributed by volunteers who identify and map roads using a variety of data sources. Reviews have shown that OSM data has a positional accuracy of 1.57 m (El-Ashmaway 2016), which was more than sufficient for our analyses because mean distance to the nearest road for carcasses and quadrature points was 3,325 m ± SD of 2,678 m. Because we were interested in how vehicle access might have influenced poaching, we excluded footpaths and bridleways from our analysis. These two road types were rare in our study area regardless. For each carcass location or quadrature point, we calculated the Euclidean distance to the nearest road in the dataset.

### Tree cover

We obtained tree cover data on a 0.00025-degree scale (∼25 x 25 m in our study area) from the Global Forest Change project, version 1.12 (Hansen et al. 2013). These data were derived from Landsat imagery and give the proportion of the cell covered by trees. We used the 2024 update, which was the most recent data available at the time of analysis. We calculated the covariate for the analysis by extracting the value for each carcass and quadrature point location.

### Human population

We obtained data on human populations from the High Resolution Population Density Maps project (Tiecke et al. 2017). This dataset uses human census data and remotely sensed building locations to estimate human populations on a 30 x 30 m grid spanning the globe.

Because our study area was sparsely populated, we aggregated data on two scales around each carcass location or quadrature point. For each point, we calculated the population within 1- and 10-km radii around the point. These distances were chose arbitrarily to represent two different levels of accessibility by local inhabitants.

### Distance to water

To measure distance to water, we used the Modified Normalized Difference Water Index (MDNWI) to map water cover in the study areas. This index has been found to be accurate for detecting waterholes in southern Africa (Dzinotizei et al. 2017). We downloaded Landsat 8 level-1 processed scenes for the study area acquired between 1 January 2023 and 15 July 2026. We removed pixels obscured by clouds or cloud shadows and then calculated MNDWI for each image and cell as:

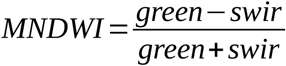

Multiple scenes were available for some portions of the study area. For each month in the dataset, we combined the available scenes and calculated the maximum extent of water cover.

Next, we divided the months into wet (November through May) and dry (June through October) seasons. For each season and year, a pixel was considered to have water if it had MNDWI ≥0 for at least one month. For each season type, pixels which had water in at least two of the four years were considered to have water for our purposes of measuring distance to water. Finally, for each season, we measured the distance between each carcass and quadrature point and the nearest cell with water.

### Distance to BDF camp

We obtained locations of BDF anti-poaching camps during aerial survey and reconnaissance flights over northern Botswana. The covariate used in the analysis was the distance to the nearest camp for each quadrature point or carcass location.

### Slope and elevation

We used Shuttle Radar Topography Mission (SRTM) data to measure elevation and slope in the study area. SRTM data provides surface elevation at a resolution of 1 arc-second. We used the "slope" function in the *raster* R package to calculate percent slope for each grid cell. We calculated the values for the analysis by extracting the slope and grid cell for whichever grid cell in which a given carcass location or quadrature point was located.

## Appendix S2. Supplemental figures

**Figure S1.**
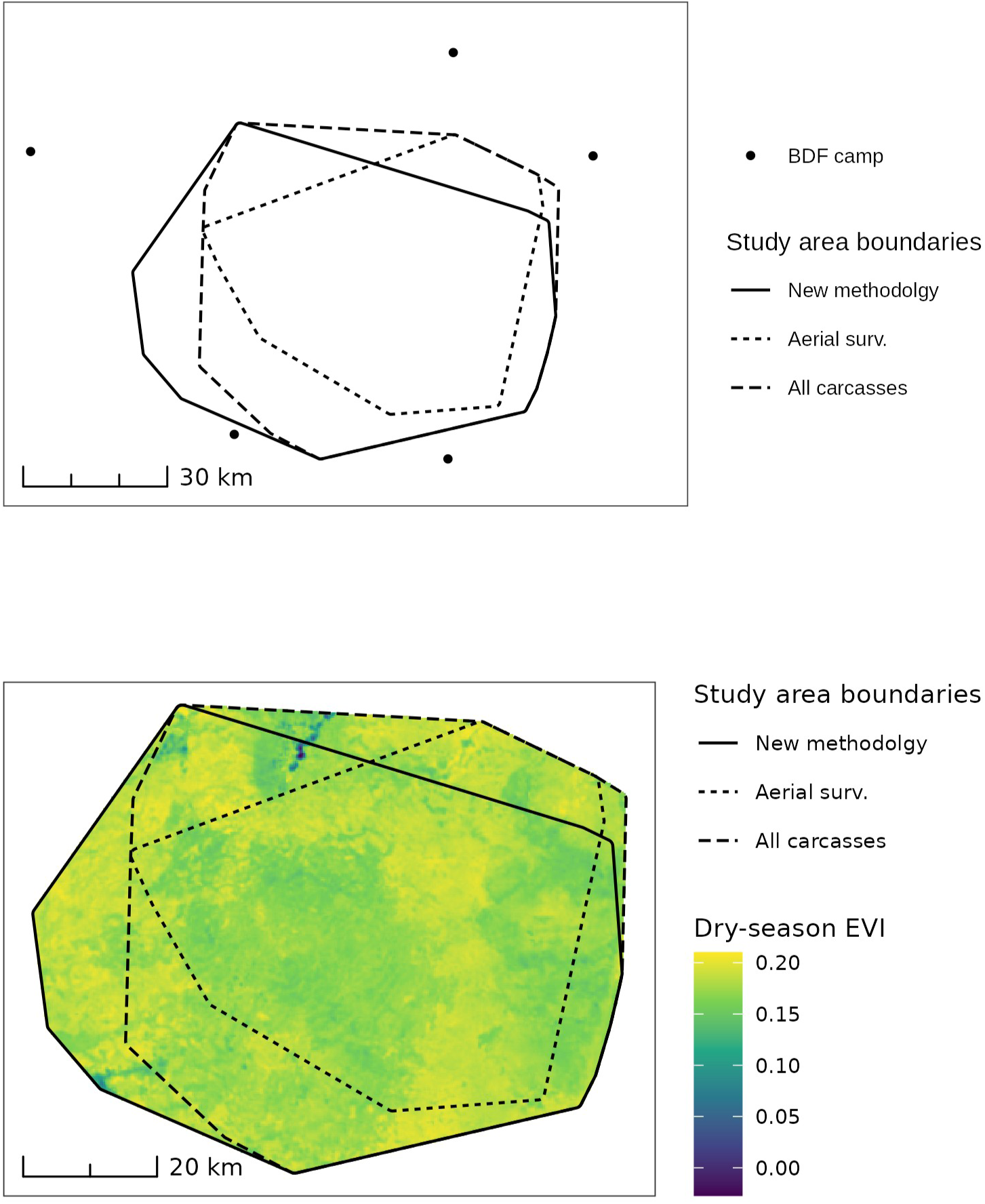
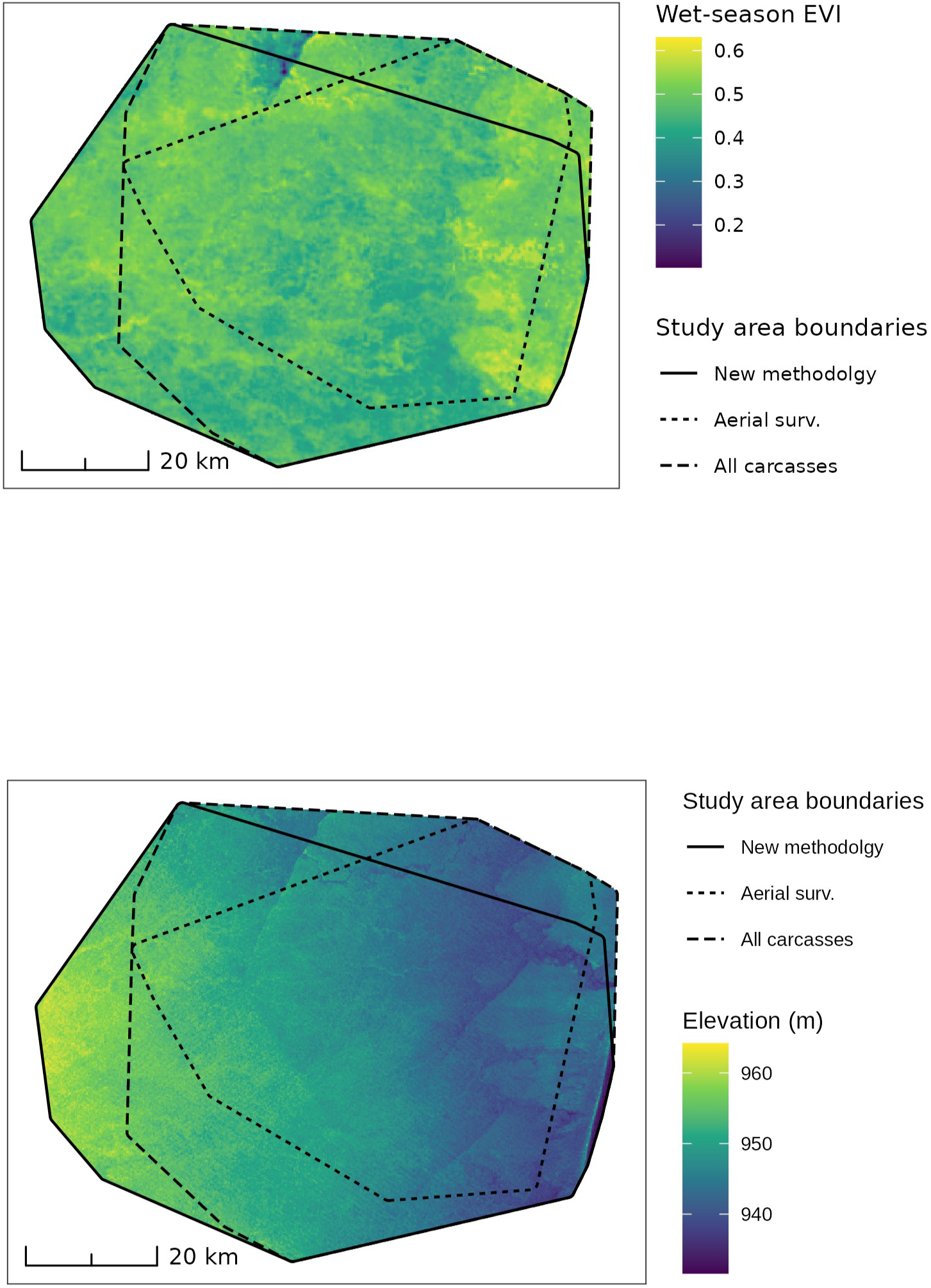
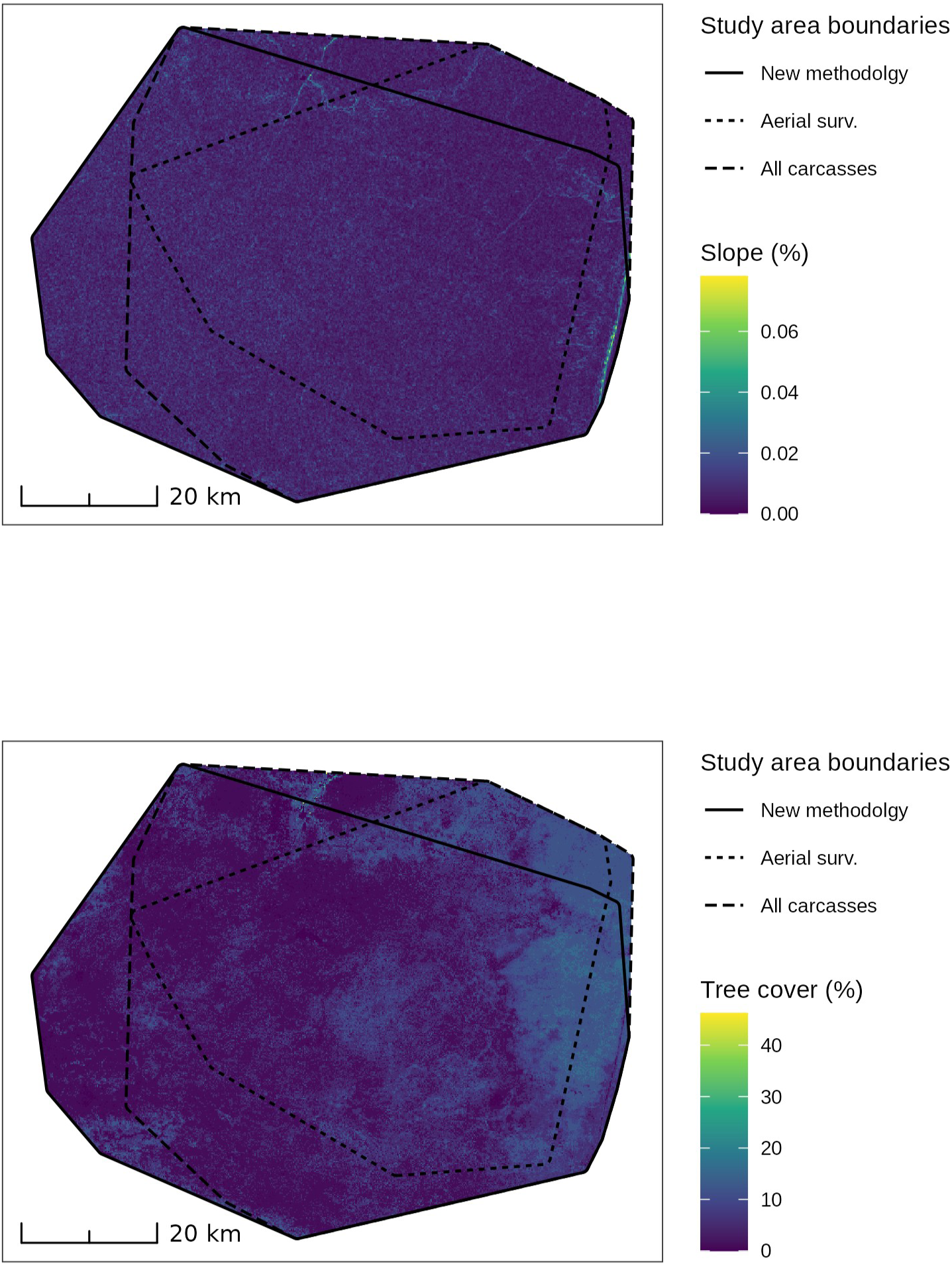
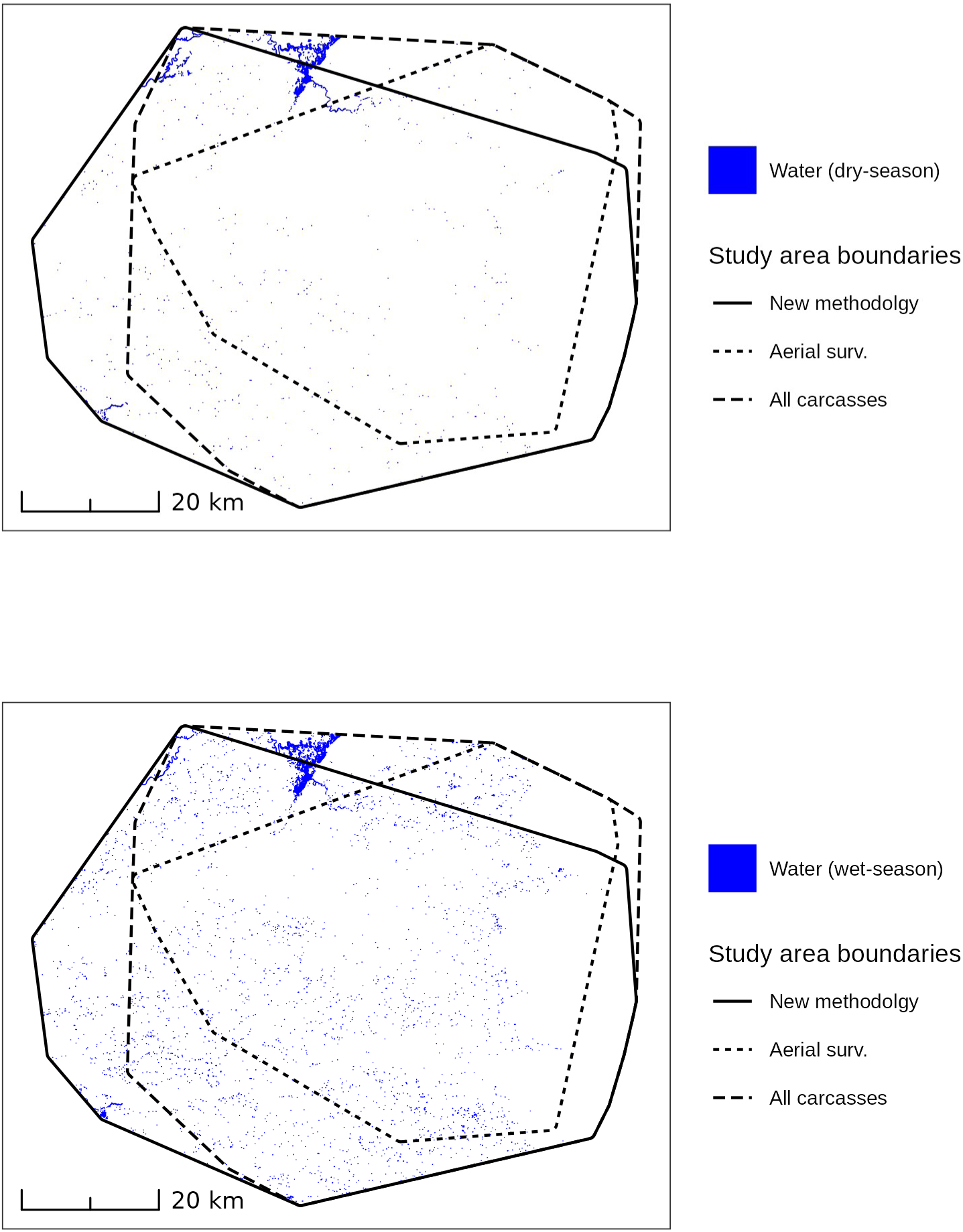
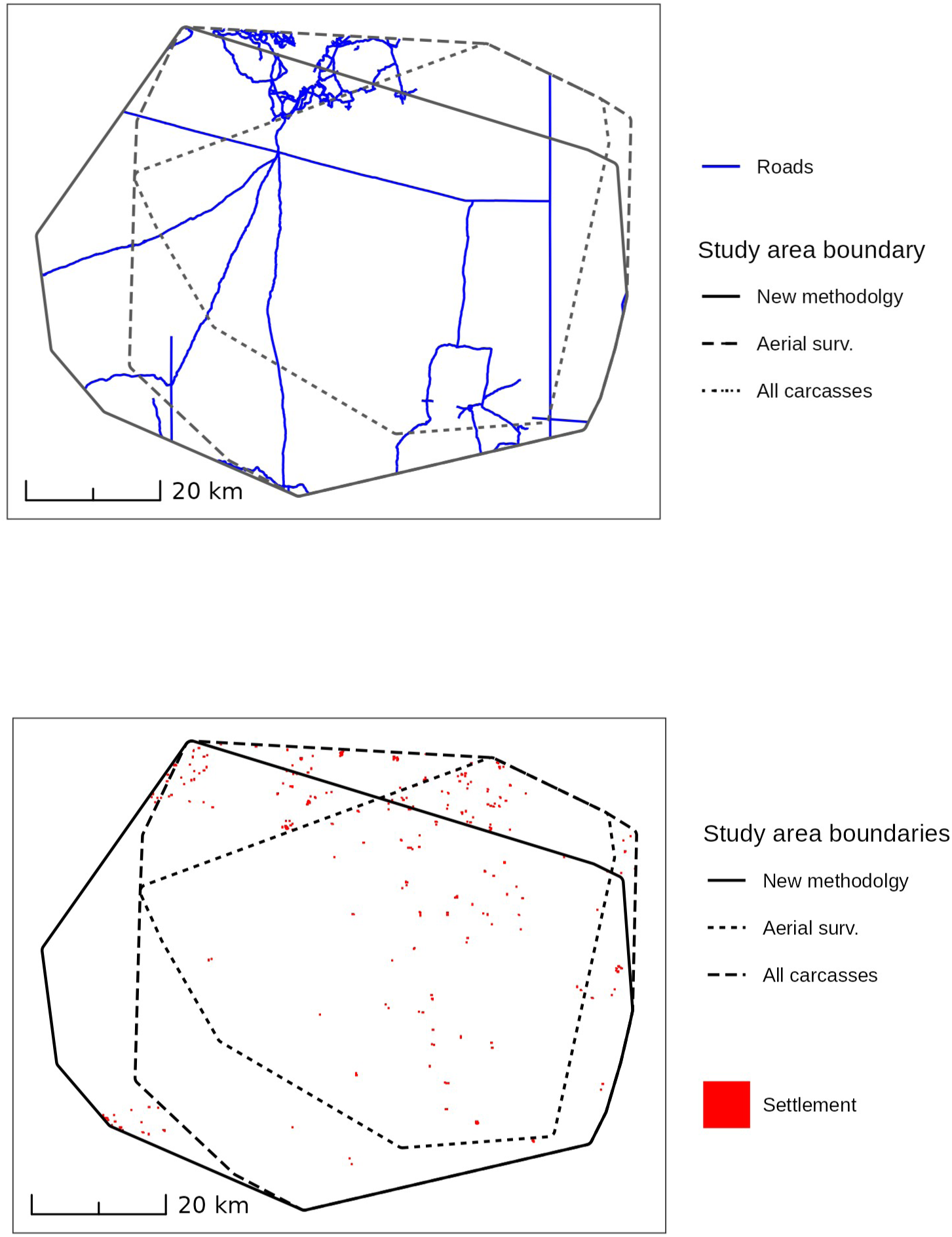
Maps of independent variables used in spatial models of elephant poaching.

**Fig. S2.**
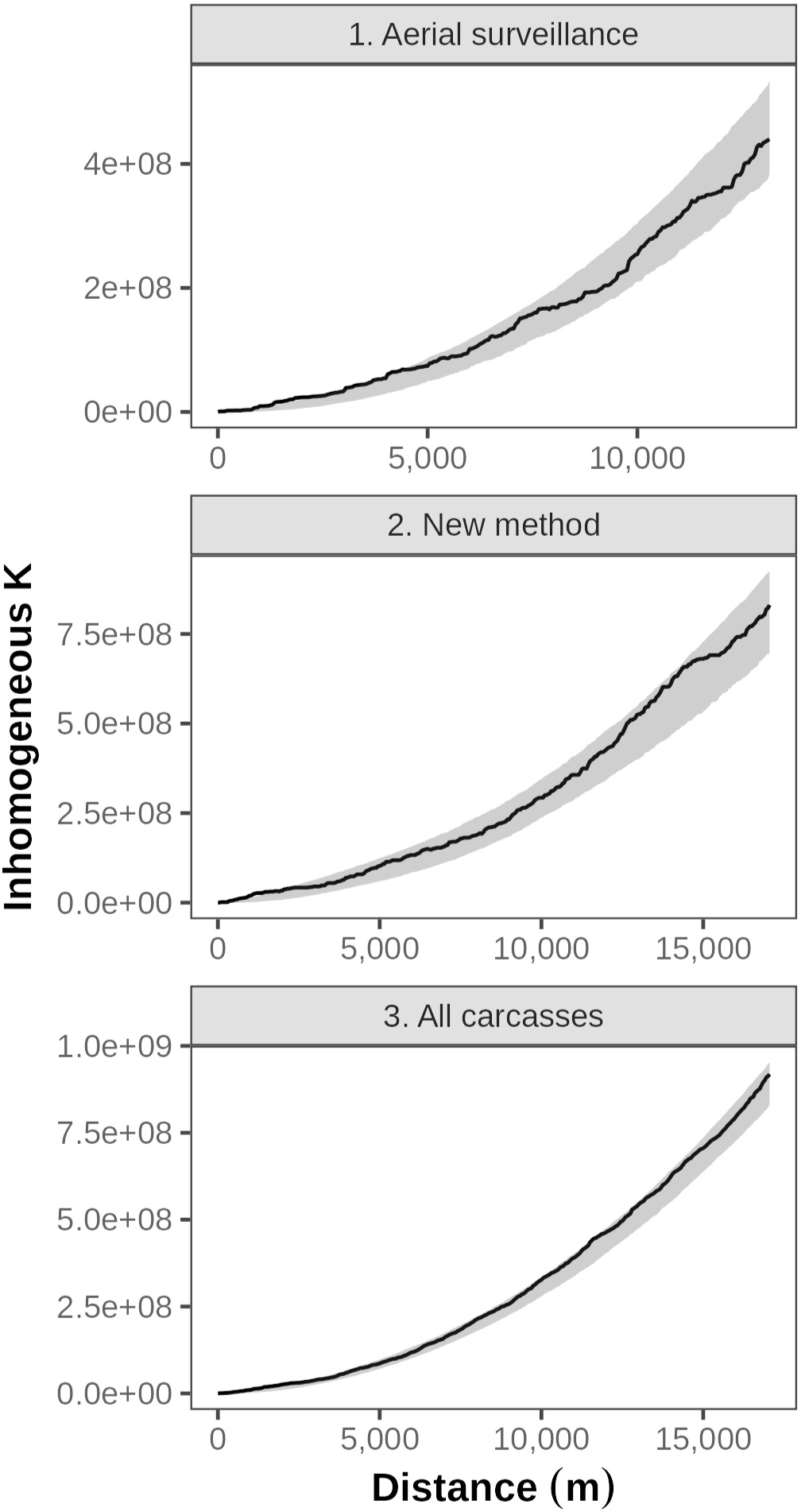
K-function plots for Poisson point process models predicting intensity of poached elephants in northern Botswana, 2023-2026. Figures are labeled by dataset

## References

Alexander K, McNutt JW, Vandewalle M (2018) Scientists question BBC reporting over elephant poaching crisis in Botswana. In: Afr. Geogr. https://africageographic.com/stories/response-recent-bbc-report-elephant-poaching-crisis-botswana/. Accessed 16 Nov 2026

Allen CRB, Brent LJN, Motsentwa T, et al (2020) Importance of old bulls: leaders and followers in collective movements of all-male groups in African savannah elephants (Loxodonta africana). Sci Rep 10:13996. 10.1038/s41598-020-70682-y

Baddeley A, Rubak E, Turner R (2015) Spatial Point Patterns: Methodology and Applications with R. Chapman and Hall/CRC, New York

Baddeley A, Turner R (2005) spatstat: An R Package for Analyzing Spatial Point Patterns. J Stat Softw 12:1–42. 10.18637/jss.v012.i06

Beale CM, Hauenstein S, Mduma S, et al (2018) Spatial analysis of aerial survey data reveals correlates of elephant carcasses within a heavily poached ecosystem. Biol Conserv 218:258–267. 10.1016/j.biocon.2017.11.016

Berman M, Turner TR (1992) Approximating Point Process Likelihoods with Glim. J R Stat Soc Ser C Appl Stat 41:31–38. 10.2307/2347614

Boyce MS, Vernier PR, Nielsen SE, Schmiegelow FKA (2002) Evaluating resource selection functions. Ecol Model 157:281–300. 10.1016/S0304-3800(02)00200-4

Bussière EMS, Potgieter D (2023a) KAZA Elephant Survey 2022, Volume I: Results and Technical Report. KAZA TFCA Secretariat, Kasane, Botswana

Bussière EMS, Potgieter D (2023b) KAZA Elephant Survey 2022, Volume II: Stratum Reports. KAZA TFCA Secretariat, Kasane, Botswana

Cassidy L, Salerno J (2020) The need for a more inclusive science of elephant conservation. Conserv Lett 13:e12717. 10.1111/conl.12717

Chase M (2011) Dry season fixed-wing aerial survey of elephants and wildlife in northern Botswana, September - November 2010. Elephants Without Borders, Kasane, Botswana

Chase M, Schlossberg, S, Landen, K, et al (2015) Dry season aerial survey of elephants and wildlife in northern Botswana, July - October 2014. Elephants Without Borders, Kasane, Botswana

Chase MJ, Schlossberg S (2019) Dry season aerial survey of elephants and wildlife in northern Botswana. Elephants Without Borders, Kasane, Botswana

Chase MJ, Schlossberg S, Griffin CR, et al (2016) Continent-wide survey reveals massive decline in African savannah elephants. PeerJ 4:e2354. 10.7717/peerj.2354

Chiyo PI, Obanda V, Korir DK (2015) Illegal tusk harvest and the decline of tusk size in the African elephant. Ecol Evol 5:5216–5229. 10.1002/ece3.1769

CITES MIKE Programme (2020) Monitoring the Illegal Killing of Elephants: Aerial Survey Standards for the MIKE Programme. Version 3.0. CITES MIKE Programme, Nairobi, Kenya

Dovers E, Stoklosa J, Warton DI (2024) Fitting Log-Gaussian Cox Processes Using Generalized Additive Model Software. Am Stat 78:418–425. 10.1080/00031305.2024.2316725

Ferreguetti ÁC, Pereira-Ribeiro J, Prevedello JA, et al (2018) One step ahead to predict potential poaching hotspots: Modeling occupancy and detectability of poachers in a neotropical rainforest. Biol Conserv 227:133–140. 10.1016/j.biocon.2018.09.009

Galeragwe M (2025) Bill seeks to allow use of arms by wildlife officers. In: Botsw. Dly. News. https://dailynews.gov.bw/news-detail/85571. Accessed 17 Nov 2026

Government of Botswana (2025) Botswana National Anti-Poaching Strategy 2025-2030. Ministry of Environment and Tourism, Gaborone, Botswana

Ibisch PL, Hoffmann MT, Kreft S, et al (2016) A global map of roadless areas and their conservation status. Science 354:1423–1427. 10.1126/science.aaf7166

Janssen J, Lemieux A, Nivette A, Ruiter S (2024) A scoping review on what motivates individuals to illegally harvest wildlife. Glob Crime 25:97–121. 10.1080/17440572.2024.2342780

Jones T, Cusack JJ, Pozo RA, et al (2018) Age structure as an indicator of poaching pressure: Insights from rapid assessments of elephant populations across space and time. Ecol Indic 88:115–125. 10.1016/j.ecolind.2018.01.030

Ketshabile L, Ferreira IW (2009) Tourism policy and the economic impact of tourism in Botswana. J Bus Manag Dyn 3:107–115

Kuiper T, Kavhu B, Ngwenya NA, et al (2020) Rangers and modellers collaborate to build and evaluate spatial models of African elephant poaching. Biol Conserv 243:108486. 10.1016/j.biocon.2020.108486

Leader-Williams N, Albon SD, Berry PSM (1990) Illegal Exploitation of Black Rhinoceros and Elephant Populations: Patterns of Decline, Law Enforcement and Patrol Effort in Luangwa Valley, Zambia. J Appl Ecol 27:1055–1087. 10.2307/2404395

Lindsay K, Chase M, Landen K, Nowak K (2017) The shared nature of Africa’s elephants. Biol Conserv 215:260–267. 10.1016/j.biocon.2017.08.021

Lupton C (2025) Illegal Wildlife Trade as a Method of Financing Terrorism. In: Goldbarsht D, de Koker L, Ferrill J (eds) The Financial War on Crime and Terrorism: Opportunities and Challenges. Springer Nature Switzerland, Cham, pp 123–144

Maingi JK, Mukeka JM, Kyale DM, Muasya RM (2012) Spatiotemporal patterns of elephant poaching in south-eastern Kenya. Wildl Res 39:234–249. 10.1071/WR11017

Marra G, Wood SN (2011) Practical variable selection for generalized additive models. Comput Stat Data Anal 55:2372–2387. 10.1016/j.csda.2011.02.004

Milner-Gulland EJ, Beddington JR (1993) The exploitation of elephants for the ivory trade: an historical perspective. Proc R Soc Lond B Biol Sci 252:29–37. 10.1098/rspb.1993.0042

Ministry of Environment and Tourism (2026) Committee of Supply Speech for Head 2000: Recurrent and Development Budget Proposals for Financial Year 2026/2027. Government of Botswana, Gaborone, Botswana

Mmolai E (2018) President Masisi clarifies reports. Botsw. Dly. News

Moller R, Cahill N, Kimaile JK (2019) The Big Tuskers of Tsavo: highlighting the importance of the Big Tusker Project in Kenya’s Tsavo Conservation Area. Int Zoo Yearb 53:189–199. 10.1111/izy.12218

Moss CJ (2001) The demography of an African elephant (Loxodonta africana) population in Amboseli, Kenya. J Zool 255:145–156. 10.1017/S0952836901001212

Muth RM, Bowe Jr. JF (1998) Illegal harvest of renewable natural resources in North America: Toward a typology of the motivations for poaching. Soc Nat Resour 11:9–24. 10.1080/08941929809381058

Ngorima P, Mpakairi KS, Kavhu B, et al (2022) Trends in elephant poaching in the Mid-Zambezi Valley, Zimbabwe: Lessons learnt and future outlook. Afr J Ecol 60:1324–1328. 10.1111/aje.13039

Paudel PK, Shrestha D, Deshar R (2025) Global patterns and drivers of wildlife hunting and trade: A review. Ecol Front 45:1147–1156. 10.1016/j.ecofro.2025.06.005

Poole JH (1989) Mate guarding, reproductive success and female choice in African elephants. Anim Behav 37:842–849. 10.1016/0003-3472(89)90068-7

Rashidi P, Wang T, Skidmore A, et al (2016) Elephant poaching risk assessed using spatial and non-spatial Bayesian models. Ecol Model 338:60–68. 10.1016/j.ecolmodel.2016.08.002

Renner IW, Elith J, Baddeley A, et al (2015) Point process models for presence-only analysis. Methods Ecol Evol 6:366–379. 10.1111/2041-210X.12352

Rija AA, Critchlow R, Thomas CD, Beale CM (2020) Global extent and drivers of mammal population declines in protected areas under illegal hunting pressure. PLOS ONE 15:e0227163. 10.1371/journal.pone.0227163

Ripple WJ, Abernethy K, Betts MG, et al (2016) Bushmeat hunting and extinction risk to the world’s mammals. R Soc Open Sci 3:160498. 10.1098/rsos.160498

Schipper J, Chanson JS, Chiozza F, et al (2008) The Status of the World’s Land and Marine Mammals: Diversity, Threat, and Knowledge. Science 322:225–230. 10.1126/science.1165115

Schlossberg S, Chase M (2024) Population trends and conservation status of elephants in Botswana and the Kavango-Zambezi Transfrontier Conservation Area. Elephants Without Borders, Kasane, Botswana

Schlossberg S, Chase MJ (in press) Trophy hunting creates a population sink for mature male elephants in Botswana. PeerJ

Schlossberg S, Chase MJ (2025) Scientific Review of Botswana’s Elephant Hunting Programme. Elephants Without Borders, Kasane, Botswana

Schlossberg S, Chase MJ, Gobush KS, et al (2020) State-space models reveal a continuing elephant poaching problem in most of Africa. Sci Rep 10:10166. 10.1038/s41598-020-66906-w

Schlossberg S, Chase MJ, Sutcliffe R (2019) Evidence of a Growing Elephant Poaching Problem in Botswana. Curr Biol 29:2222–2228. 10.1016/j.cub.2019.05.061

Shaffer MJ, Bishop JA (2016) Predicting and Preventing Elephant Poaching Incidents through Statistical Analysis, GIS-Based Risk Analysis, and Aerial Surveillance Flight Path Modeling. Trop Conserv Sci 9:525–548. 10.1177/194008291600900127

Slotow R, van Dyk G, Poole J, et al (2000) Older bull elephants control young males. Nature 408:425–426. 10.1038/35044191

Thouless CR, Dublin HT, Blanc JJ, et al (2016) African Elephant Status Report 2016: an update from the African Elephant Database. IUCN, Gland, Switzerland

Wang S, Tian Z (2024) New challenges in financial supervision: environmental crime terrorism financing. Trends Organ Crime 27:212–228. 10.1007/s12117-023-09500-6

Wasser SK, Brown L, Mailand C, et al (2015) Genetic assignment of large seizures of elephant ivory reveals Africa’s major poaching hotspots. Science 349:84–87. 10.1126/science.aaa2457

Wittemyer G, Daballen D, Douglas-Hamilton I (2013) Comparative Demography of an At-Risk African Elephant Population. PLoS ONE 8:e53726. 10.1371/journal.pone.0053726

Wood S (2017) Generalized Additive Models: an Introduction with R, 2nd edn. CRC press

Zhou X, Wang Q, Zhang W, et al (2018) Elephant poaching and the ivory trade: The impact of demand reduction and enforcement efforts by China from 2005 – 2017. Glob Ecol Conserv 16:e00486. 10.1016/j.gecco.2018.e00486

## References

Dzinotizei Z, Murwira A, Zengeya FM, Guerrini L (2018) Mapping waterholes and testing for aridity using a remote sensing water index in a southern African semi-arid wildlife area. Geocarto International 33:1268–1280. 10.1080/10106049.2017.1343394

El-Ashmawy KLA (2016) Testing the positional accuracy of OpenStreetMap data for mapping applications. Geodesy and Cartography 42:25–30. 10.3846/20296991.2016.1160493

Tiecke TG, Liu X, Zhang A, et al (2017) Mapping the world population one building at a time. Arxiv.org. 10.48550/arXiv.1712.05839

